# Defects in the assembly of ribosomes selected for β-amino acid incorporation

**DOI:** 10.1101/733584

**Authors:** Fred R. Ward, Zoe L. Watson, Omer Ad, Alanna Schepartz, Jamie H. D. Cate

**Affiliations:** Department of Molecular and Cell Biology, University of California-Berkeley, Berkeley, CA; Department of Chemistry, University of California-Berkeley, Berkeley, CA; Department of Chemistry, Yale University, New Haven, CT; Molecular Biophysics and Integrated Bioimaging Division, Lawrence Berkeley National Laboratory, Berkeley, CA

**Keywords:** ribosome, ribosome structure, peptidyl transferase center, beta-amino acids, ribosome assembly

## Abstract

Ribosome engineering has emerged as a promising field in synthetic biology, particularly concerning the production of new sequence-defined polymers. Mutant ribosomes have been developed that improve the incorporation of several non-standard monomers including D-amino acids, dipeptides, and β-amino acids into polypeptide chains. However, there remains little mechanistic understanding of how these ribosomes catalyze incorporation of these new substrates. Here we probed the properties of a mutant ribosome–P7A7–evolved for better *in vivo* β-amino acid incorporation through *in vitro* biochemistry and cryo-electron microscopy. Although P7A7 is a functional ribosome *in vivo*, it is inactive *in vitro*, and assembles poorly into 70S complexes. Structural characterization revealed large regions of disorder in the peptidyltransferase center and nearby features, suggesting a defect in assembly. Comparison of RNA helix and ribosomal protein occupancy with other assembly intermediates revealed that P7A7 is stalled at a late stage in ribosome assembly, explaining its weak activity. These results highlight the importance of ensuring efficient ribosome assembly during ribosome engineering towards new catalytic abilities.

## Introduction

The ribosome is a large molecular machine capable of directing the polymerization of a distinct sequence of amino acids by decoding a messenger RNA (mRNA) template. Its ability to accurately and efficiently select and incorporate the correct monomer from a pool of over 20 substrates makes the ribosome one the most versatile machines for polymer synthesis. No other known methods, from solid-state chemistry to bulk polymerization, can achieve both the specificity and yield of ribosome-catalyzed synthesis, particularly for products longer than ~50 monomers in length.^1^ The ribosome has thus become a target of engineering efforts to eventually develop a ribosome-based platform for the synthesis of new classes of sequence-defined polymers. The large obstacle to these engineering attempts is the several-billion-year optimization of the ribosome, and the rest of the translational apparatus, toward efficient and selective α-amino acid polymerization. New sequence-defined polymerization chemistries will likely require repurposing a suite of enzymes and nucleic acids, including tRNAs,^2,3^ aminoacyl tRNA synthetases (aaRSs),^4,5^ elongation factor Tu (EF-Tu),^6,7^ in addition to the ribosome itself.^8-12^

The catalytic core of the ribosome, the peptidyl transferase center (PTC), induces the attack of an A-site aminoacyl-tRNA on the ester bond of the P-site peptidyl-tRNA, transferring and extending the growing polypeptide chain. The PTC is thought to encourage proper polymerization through several mechanisms including orienting the substrates,^13^ favoring productive acid-base chemistry,^14^ and protecting the peptidyl-tRNA from dead-end hydrolysis.^15^ Like many other enzymes, the PTC undergoes a substantial activating rearrangement upon substrate binding that can accommodate a wide range of monomer side chains. However, the ribosome only weakly catalyzes polymerization of substrates with different backbone chemistries, including peptoids,^16^ N-methyl amino acids,^17–19^ D-amino acids,^20^ aromatic foldamers,^21,22^ aramids,^23^ malonates,^23^ and β^3^-amino acids.^24^

β^3^-amino acids (referred to here as β-amino acids) are a useful model substrate for novel ribosome polymerization reactions as they introduce new challenges for catalysis while closely resembling the natural α-amino substrates. The extra methylene between the alpha carbon and carboxylic acid groups adds more bulk, rotational degrees of freedom, and diminishes reactivity,^25^ but the nucleophile remains an amino group, as in α-amino acids. The extra methylene group confers advantages to β-amino acid oligomers as peptidomimetics, as they form stable folds, display reduced immunogenicity, and are more resistant to proteolysis.^26–28^ Importantly, despite reacting poorly in the PTC, under certain conditions β-amino acids can be consecutively polymerized by the ribosome *in vitro*.^3^ Their similarities to natural monomers have made β-amino acids an attractive substrate target for ribosome engineering.

Attempts to engineer new polymerization chemistry into the ribosome have understandably focused on mutations to the PTC. Aggressive mutations of 10+ bases in the PTC have generated ribosomes that can better translate D-amino acids,^12^ dipeptide substrates,^29^ polyproline motifs,^8^ and β-amino acids.^9,10^ These mutations have been proposed to solve the space, orientation, and reactivity problems inherent to many backbone-modified substrates, but no mechanistic studies of these improved ribosomes have been published. Indeed, most of these mutant ribosomes have only been tested *in vivo* or in lysate-based cell free systems with wild-type ribosomes present, which allows subunit exchange. Although these ribosomes represent exciting steps toward new sequence-defined polymerization capabilities, a detailed understanding of how PTC mutations help widen the substrate scope of the ribosome is still lacking. In particular, the PTC is highly conserved across all domains of life and large-scale mutations may have knock-on effects that limit ribosome utility. Therefore, these engineering problems must always be evaluated with a discerning focus on the mechanistic implications of such changes, in order to better understand how these PTC alterations affect ribosome activity.

Here were present structural and biochemical characterization of a β-amino acid translating ribosome, P7A7, discovered via an *in vivo* selection for β-puromycin incorporation.^9^ The P7A7 PTC carries twelve mutations over two regions near the tRNA A site and exit tunnel. We show that purified P7A7 ribosomes are inactive during *in vitro* translation and do not form stable 70S complexes. A cryo-electron microscopy (cryo-EM) structure of P7A7 50S ribosomal subunits reveals substantial disordering of the PTC and nearby inter-subunit bridge helices when compared with an equivalent wild-type (WT) 50S structure. Analysis of the P7A7 map reveals a depletion of late-assembling ribosomal proteins, which was confirmed using tandem mass tag (TMT) relative quantitation. When compared with existing studies of 50S ribosomal subunit assembly, P7A7 appears trapped as a late assembly intermediate, explaining its poor activity. Our results suggest that the radical PTC mutations often seen in engineered ribosomes may have unintended effects on ribosome assembly and stability that limit the utility of many of these variants.

## Results

Prior analysis of the P7A7 ribosome and its precursor, 040329, was limited to *in vivo* or lysate based characterization in the presence of wild-type ribosomes.^9,10^ This natural background translation makes it challenging to accurately understand P7A7’s altered substrate scope, necessitating the development of a pure P7A7 translation system. Initial attempts at creating *rrn* knockout ‘Squires’ strains carrying a P7A7-23S rRNA encoding variant of the *rrnb* operon on a plasmid failed, indicating that P7A7 cannot support cellular growth on its own.^30^ While the 12 PTC mutations of P7A7 likely diminish its efficiency at natural protein synthesis, it had been observed to synthesize full-length dihydrofolate reductase *in vivo*, suggesting a level of translational competency amenable to study. We assayed the levels of P7A7 50S ribosomal subunits in polysome fractions from mid-log phase *E. coli* Mach1 cells using a semi-quantitative reverse transcription-PCR (RT-PCR) assay, recognizing an MS2 RNA tag grafted onto helix 98 of the 23S rRNA.^31^ P7A7 ribosomes are depleted, but not absent, in both 70S and polysome fractions **(Figure 1A; Figure S1A),** indicating that P7A7 50S subunits are deficient in subunit association. To remove background WT ribosomes and develop an isolated P7A7 translation system, we purified P7A7 50S subunits using affinity chromatography of MBP-phage MS2 coat protein fusion bound to a tag on helix 98 **(Figure S1B).**

**Figure 1.**
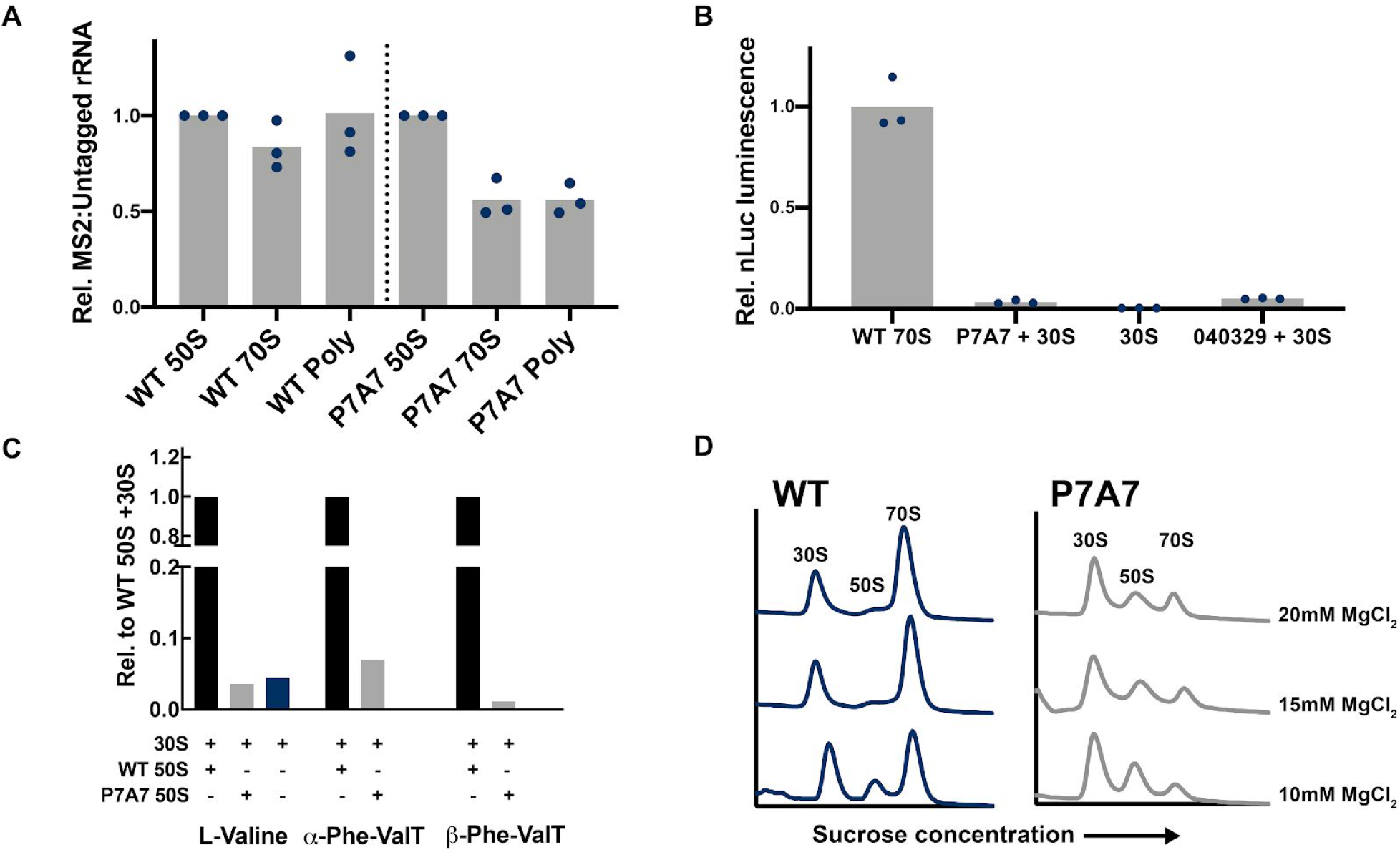
WT and P7A7 ribosome activity. A) WT-MS2:WT or P7A7-MS2:WT rRNA ratios in different polysome profile fractions, normalized to 50S ratio per replicate. Relative quantities of rRNA measured by RT-PCR over the MS2-tag region and quantification of bands resolved by polyacrylamide gel electrophoresis. Bars represent mean values. B) Nanoluciferase expression measured from PURExpress Δ-ribosome *in vitro* translation reactions with different purified ribosomes. Values reported relative to WT 70S control. Bars represent means. C) LC-MS monitored MXFDYKDDDDK peptide synthesis from PURExpress Δ-ribosome-tRNA-amino-acid *in vitro* translation reactions with varying purified ribosomes. The tRNA for position X was either Val-tRNA_val_, Phe-tRNA_val_ (flexizyme charged), or β^3^-Phe-tRNA_val_ (flexizyme charged). Single replicates are shown for each condition. D) *in vitro* apo-ribosome association at differing Mg^2+^ concentrations, measured by sucrose gradient centrifugation and fractionation for WT 50S + WT 30S and P7A7 50S + WT 30S ribosomes. Representative gradients are shown.

We tested the activity of purified P7A7 50S subunits with WT 30S subunits in the PURExpress delta-ribosome *in vitro* transcription-translation system.^32^ Translation of a standard α-amino acid nanoluciferase reporter with P7A7 was barely discernible from background WT contamination **(Figure 1B).** Similar results were found with the earlier-generation 040329 ribosome **(Figure 1B**). While *in vitro* translation is known to be far less efficient than seen *in vivo*, the minute activity of purified P7A7 ribosomes was unexpected in light of P7A7’s DHFR synthesis^9^ and polysome occupancy *in vivo*. After additional purification of P7A7 50S subunits to remove WT contamination, short FLAG-derived α- and β-amino acid containing peptides could not be detected above background levels **(Figure 1C; Figure S2).** Because subunit association is a necessary step during translation initiation, we compared the magnesium-dependent subunit association of P7A7 and WT ribosomes. Apo-P7A7 ribosomes require high concentrations of magnesium to form 70S complexes, suggesting a defect in association that could impair translation **(Figure 1D).**

To better understand the basis for P7A7’s low activity, we solved the cryo-EM structure of its large subunit, alongside a WT 50S subunit as a control. Images were collected on a 300-kV Titan Krios microscope with a K2 direct electron detector, and processed with RELION^33^ and cryoSPARC.^34^ Global resolutions achieved for the two maps were 3.11 Å and 3.20 Å, respectively **(Figure S3).** Intriguingly, although both maps have substantial regions of the 50S subunit at high resolution, large portions of the ribosome that are resolved in the WT structure are absent in the P7A7 map **(Figure 2)**. Heterogeneous ab initio reconstructions of the P7A7 50S subunit revealed two classes of particles that represent the large subunit with two different types of disorder. The major class lacks density for ribosomal proteins uL6, uL10, uL11, uL16, bL33, and bL35 (excluding proteins bL31 and bL36 which are known to disassociate during ribosome isolation),^35^ as well as multiple stretches of the 23S rRNA comprising the PTC, helices H44, H71, H89-H93, and the end of H39. The smaller class further lacks the central protuberance entirely **(Figure S4** for workflow).

**Figure 2.**
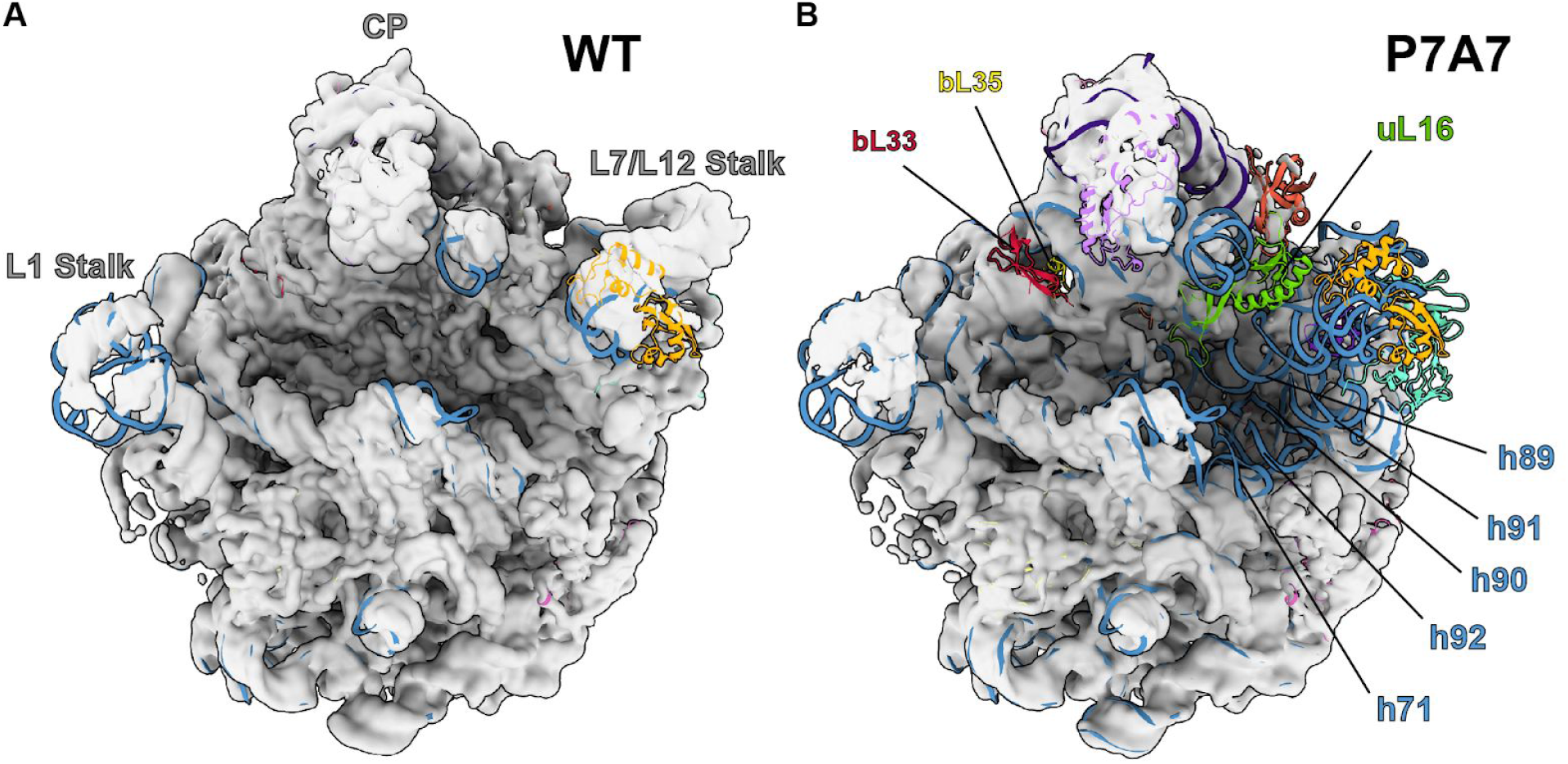
Comparison of WT 50S (A) and P7A7 (B) large subunit cryo-EM maps. Maps are low-pass filtered to 8 Å for clarity of structural features, and 50S coordinates from PDB 4YBB^38^ are docked to identify missing components. The central protuberance (CP), L1 Stalk, and L7/L12 Stalk are identified on the WT structure, and features confirmed to be missing from P7A7 by combined cryo-EM and LC-MS/MS analysis are labeled on the P7A7 structure.

Because cryo-EM often cannot resolve whether features of the ribosome are missing due to their absence from the complex or due to flexibility, we also performed relative quantitation of ribosomal proteins (RPs) using liquid chromatography with tandem mass tag (TMT) spectrometry (LC-MS/MS), with TMT labeling on WT and P7A7 50S subunits.^36^ Normalized ratios of P7A7:WT RP levels generally matched the structural observations. P7A7 is depleted of RPs uL16, bL33, and bL35 **(Figure 3).** We were able to detect stoichiometric levels of RPs uL6, uL10, and uL11, whose cryo-EM density is often weak due to motion in the large subunit arms.^35,37^

**Figure 3.**
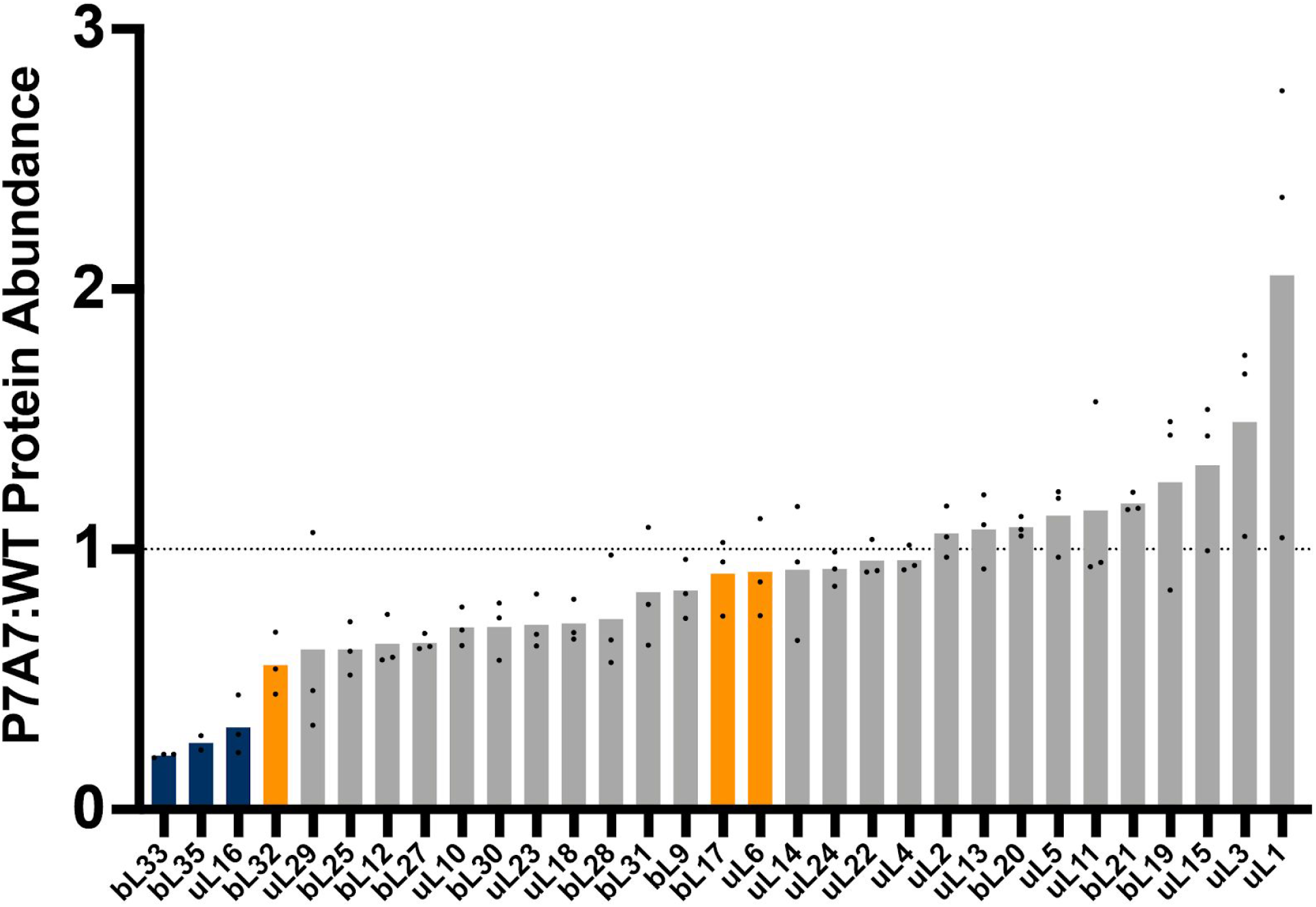
50S ribosomal protein ratios detected by LC-MS/MS and TMT relative quantitation between P7A7 and WT ribosomes. Bars represent means and are colored by absence in final assembly states (E4 and E5) of Davis and Tan, *et al.* (orange) or absence in both Davis and Tan, *et al.* and Nikolay, *et al.* states (IV and V) (blue).^35,37^

In the vicinity of the P7A7 mutations in the PTC, we observe a stark contrast between well-ordered regions seen in the WT structure, with features like base stacking clearly resolved, and adjacent regions where the density is mainly noise **(Figure 4).** Comparison of these disordered regions in the P7A7 50S subunit map with the corresponding region in the WT PTC shows several interactions that would be perturbed by some of the mutations **(Figure 5),**^38^ and may be responsible for the structural defects we observe. In particular, two base triples–the first between C2063, A2450, and C2501, and the second between C2507, G2553, and G2582− would not be able to form because of the C-A and C-U mutations at positions 2063 and 2507, respectively. Other disrupted interactions include the G2057-C2611 and G2505-C2610 base pairs **(Figure 5).**

**Figure 4.**
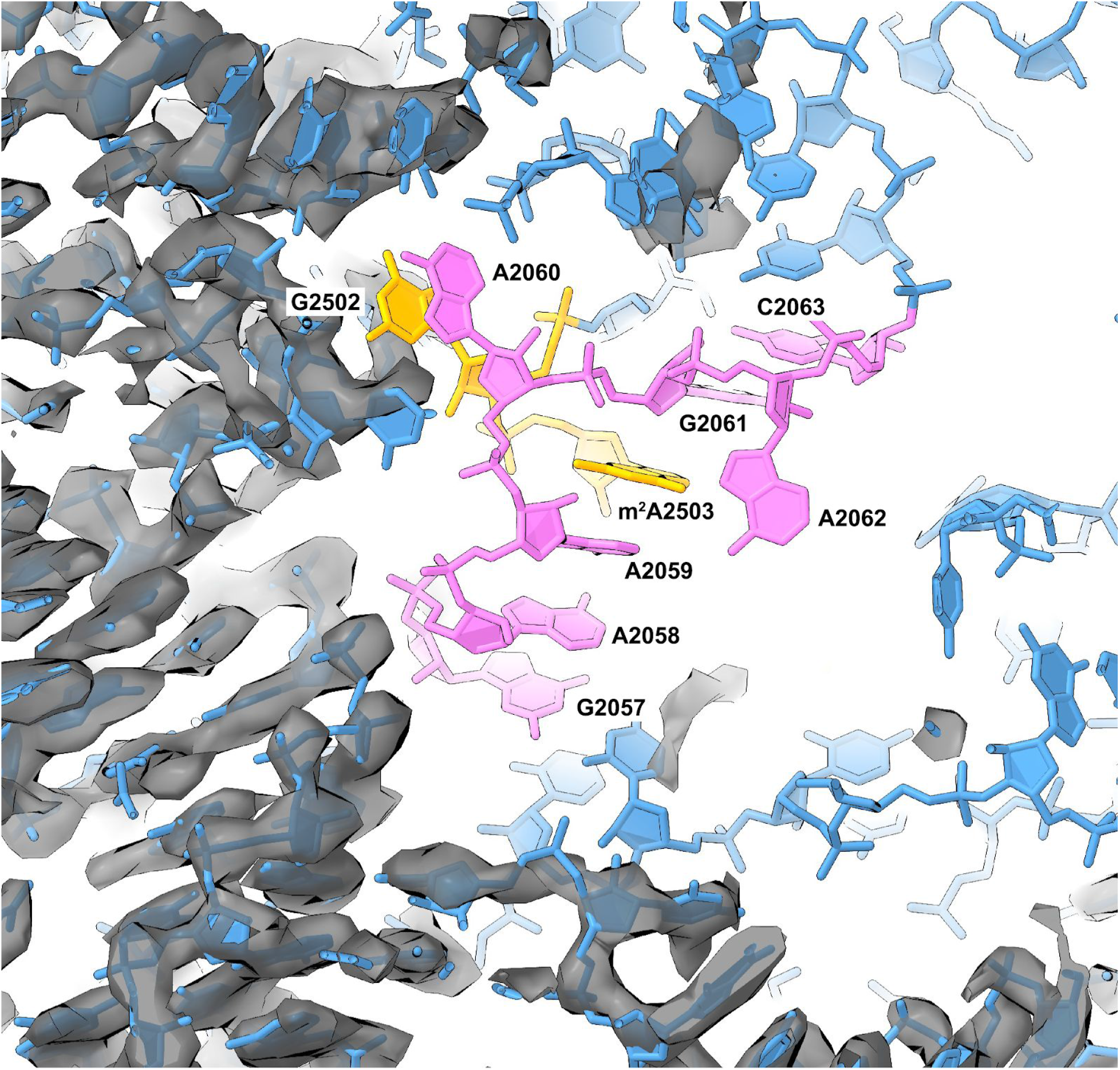
Disorder in the PTC of P7A7 50S subunits. Model of the wild-type 23S rRNA in the vicinity of the PTC superimposed on the map of the P7A7 50S subunit. Regions that are well ordered include helices H26, H35, and several loops from domain II. Mutated PTC residues 2057-2063 and 2502-2503 in P7A7 are depicted in pink and gold, respectively. The “hide dust” feature in ChimeraX^73^ was used to remove noise for clarity.

**Figure 5.**
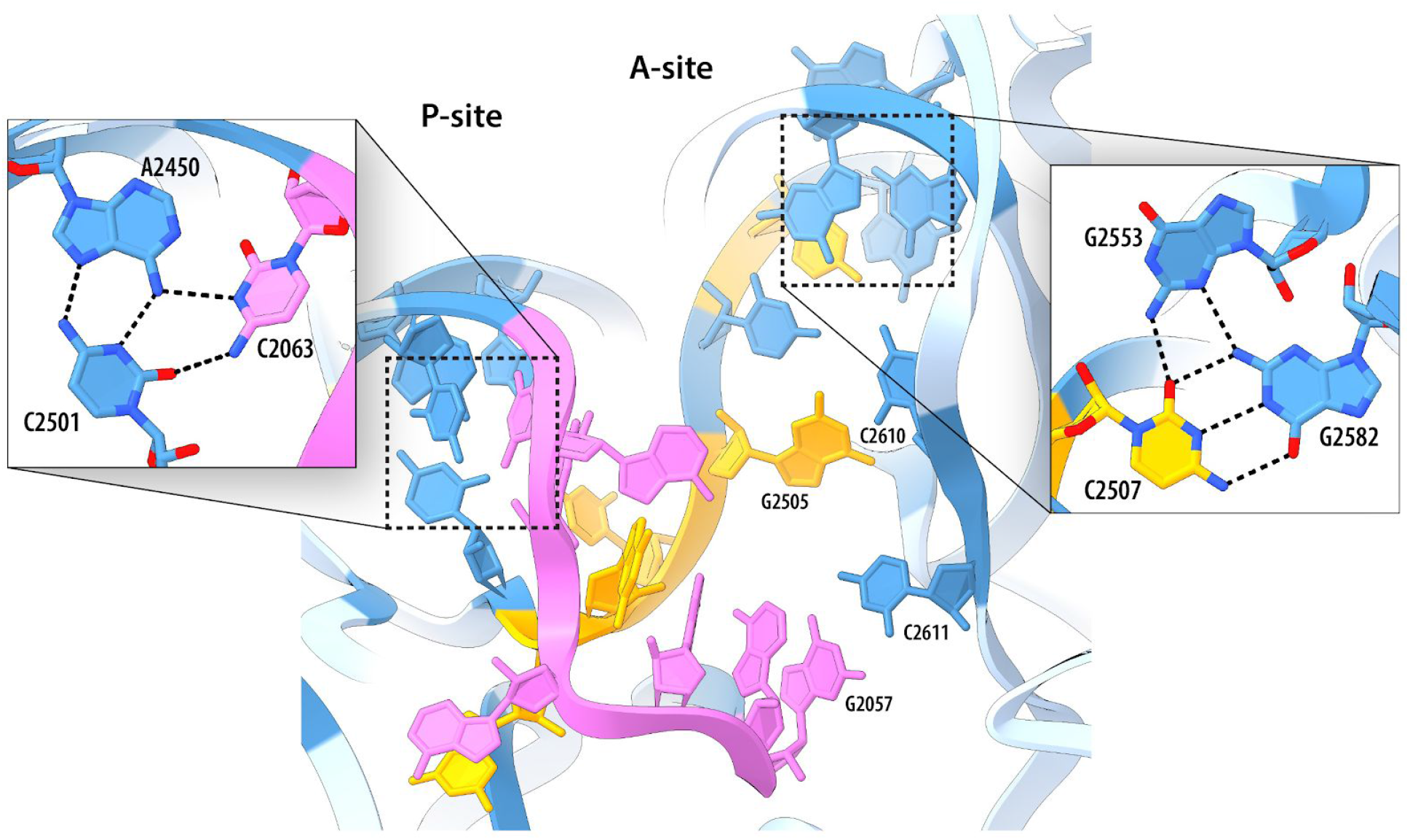
WT 23S rRNA residues directly affected by P7A7 mutations. Coordinates shown are from PDB 4YBB.^38^ Mutated residues 2057-2063 and 2502-2505, 2507 in P7A7 are depicted in pink and gold, respectively. Unaltered bases within 3 Å of mutations are shown in darker blue, and those further away are lighter blue. Close-ups highlight base triples that would be disrupted by mutations, and two disrupted base pairs are also labeled.

Low-pass filtering of the P7A7 50S subunit map allowed for some interpretation of broader features of its structure that show poor connectivity and are more difficult to discern in the high resolution map. Notably, there appears to be some helical RNA density spanning the PTC, which does not fit with the WT model **(Figure 6).** This connectivity is strikingly similar to the non-native PTC organization seen in late intermediates of the 50S assembly pathway both *in vivo* and *vitro*.^35,37^ While ribosome biogenesis is a complex process, a general order of rRNA helix formation and ribosomal protein association is known.^39,40^ Despite multiple assembly routes, resulting in a spectrum of assembly intermediates, he PTC is usually observed as the last-assembling 50S motif.

**Figure 6.**
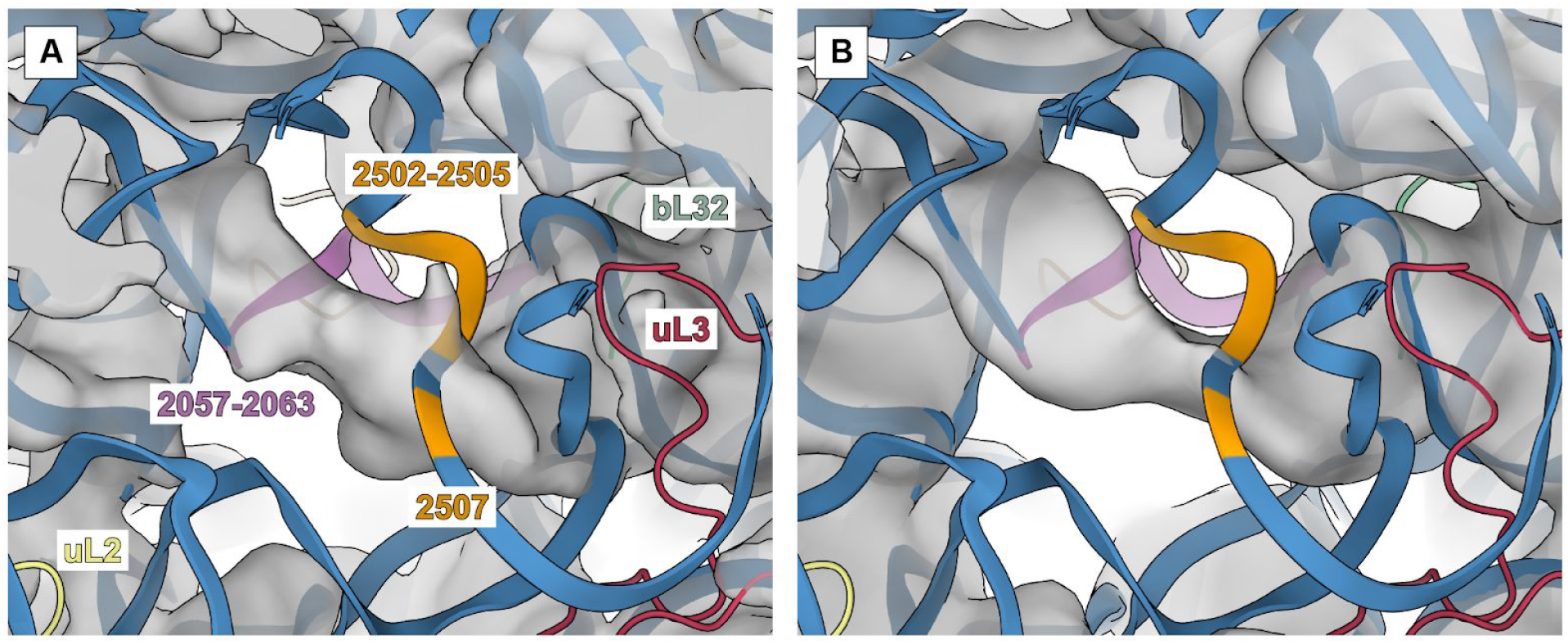
Comparison of PTC in the P7A7 map (A) and an assembly intermediate, E4, determined by Davis and Tan, *et al.* (B). Both maps show helical density that does not appear in fully assembled WT structures. P7A7 mutations regions 2057-2063 and 2502-2505, 2507 are colored in pink and orange, respectively. Other 23S rRNA regions are in blue, and nearby proteins uL2, uL3, and bL32 are yellow, red, and green, respectively.

To further explore the relationship of the observed P7A7 structure with assembly pathways, we carried out a detailed tabulation of structural features across the P7A7 and WT 50S subunit EM maps and compared them to known assembly intermediates, including states IV and V from characterization of *in vitro* reconstitution^35^ and states E4 and E5 from *in vivo* intermediates caused by bL17 knockdown.^37^ Using a previously developed calculation method^37^ we generated a comprehensive list of the cryo-EM occupancies for every ribosomal protein and rRNA helix across all 6 structures **(Figure 7A).** Hierarchical clustering grouped P7A7 alongside other PTC-unfolded states, with P7A7 most closely aligned to E4. However, ribosomal protein occupancy, particularly the absence of bL17 and bL32 from E4, revealed P7A7 as a distinct intermediate. Our TMT quantitation results **(Figure 3)** correlate well with these analyses. Nevertheless, the large-scale organization of several intermediates is quite similar despite the different methods of assembly perturbation, suggesting that a PTC folding defect is a common roadblock in 50S ribosomal subunit maturation. Interestingly, careful analysis of P7A7’s migration on a sucrose gradient reveals that it sediments slower than mature WT 50S subunits, similar to the immature 48S particle seen during *in vitro* reconstitution experiments^35^ **(Figure 7B).** Together, our structure, mass spectrometry, and biochemical results suggests that P7A7’s minimal activity *in vitro* stems primarily from errors in assembly, leaving the ribosome without a mature PTC.

**Figure 7.**
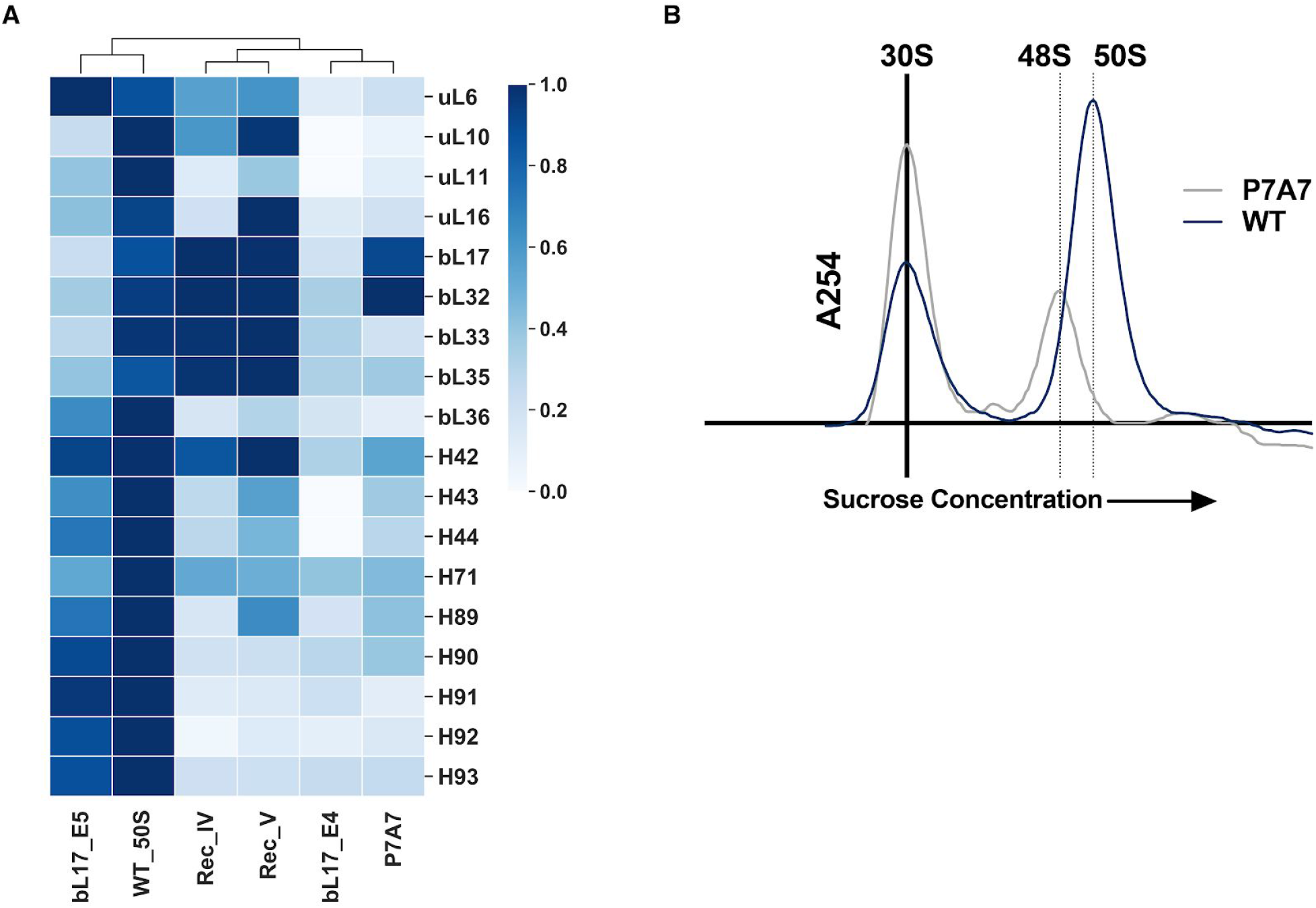
Analysis of P7A7 Assembly. A) Heatmap of 50S subunit rRNA helices and protein occupancy across cryo-EM structures of several assembly intermediates, using the method of Davis and Tan, *et al.*^37^ The heatmap is hierarchically clustered across assembly intermediates. Universally present features have been removed for clarity. B) Sucrose gradient fractionation of MS2-tag purified WT and P7A7 50S subunits run with WT 30S subunits as a standard. The 254 nm absorbance traces are shown, and the estimated sedimentation coefficient for P7A7 is marked.

## Discussion

Ribosome engineering holds promise to expand efficient template-directed polymer synthesis beyond traditional peptide chemistry, unlocking entirely new classes of materials. Toward this end, there have been several attempts at mutating the ribosome PTC to catalyze the synthesis of difficult sequences or accept new substrates, demonstrating some level of success in expanding the chemistry allowed by the ribosome. Many of these ribosomes, including P7A7 studied here, carry a large number of mutations to highly conserved bases in the PTC and nearby exit tunnel. These base mutations are necessary to overcome the strong optimization of the WT PTC toward natural protein synthesis. However, conservation of nucleotides within the PTC is likely not for catalytic ability alone. Ribosome assembly, and particularly the folding of the protein-sparse PTC, is also dependent on rRNA sequence, presenting a multi-parameter optimization problem when engineering ribosomes. While there is evidence that the ribosome possesses an innate tolerance for mutations to the PTC, even to highly conserved bases,^41^ a dozen or more mutations exceeds this plasticity. Due to these constraints, many engineered ribosomes likely possess assembly defects that limit their functionality, even if they perform better than WT ribosomes in certain challenging polymerization reactions. Further confounding engineering efforts, RNA in general shows less structural stability *in vitro*,^42^ possibly exacerbating assembly defects to the point of complete inactivity. These challenges have prevented detailed study of altered-substrate ribosomes *in vitro*, an important step in understanding how new polymerization chemistries are accessed in a remodeled PTC.

There are many plausible hypotheses that explain the effect of PTC mutations on ribosome substrate specificity, including opening up space for bulkier groups (as with β-amino acids and dipeptide substrates), properly orienting flexible substrates (β-amino acids), and rearranging suboptimal nucleophiles (i.e. poly-proline stretches, D-amino acids). These mechanistic questions will remain challenging without interpretable structures of these engineered ribosomes and a detailed understanding of conformational coupling of ribosome activities. The general disorder of P7A7’s PTC (**Figure 4**), in contrast with its *in vivo* activity (Figure 1A), suggests that disruption of rigid structures in the PTC may be a contributing mechanism to its acceptance of β-amino acids. It is possible that the PTC of P7A7 is more tolerant of a larger β-substrate only because its flexibility allows an accommodating rearrangement. Analogous promiscuity has been observed in protein enzymes, where flexibility in the active site is correlated with wider substrate specificity.^43–45^ Furthermore, the ribosome is often described as an “entropy trap,” in that proper substrate positioning is the primary contributor to highly efficient catalysis.^13^ The main barrier to β-amino acid polymerization on the ribosome would then be due to improper geometric constraints for β-substrates and not their altered reactivity. An induced-fit mechanism of PTC catalysis has been proposed to be the source of the ribosome’s side-chain promiscuity,^15,46^ albeit with less disorder than observed in P7A7. As the mutations in P7A7 lead to a less ordered, more flexible PTC fold, we can propose a model in which the additional flexibility confers additional substrate promiscuity. However, this comes at a large cost to overall catalytic efficiency.

A major contributing factor towards P7A7’s inactivity is the differential RNA structural stability often seen when comparing *in vitro* environments and the cellular milieu. Among the chemical and physical differences between these environments, crowding and divalent salt concentration are commonly implicated for their large effects on RNA structure.^42^ Success in improving conditions for RNA folding by mimicking a crowded macromolecular environment (up to 300 mg/mL^47^) provides evidence for the importance of excluded volume effects.^48–50^ Notably, *in vitro* translation reactions have been improved by optimizing molecular crowding, reducing agents, and ionic concentrations.^51–53^ Nevertheless, even these reconstituted in vitro translation systems are several orders of magnitude less complex than the bacterial cytosol. Missing are many small molecules and proteins that can interact with RNA to ensure functional folding. For example high magnesium concentrations are often used to stabilize RNA conformations *in vitro*, presumably acting as a stand-in for missing parts of the cellular environment. However, *in vivo* RNA structure probing has shown that magnesium can be a poor substitute.^50^

Stable RNA folding is particularly important for the PTC, which has a high RNA:protein ratio and folds late in the 50S subunit maturation process.^54–56^ With no ribosomal proteins to chaperone or stabilize assembly, the PTC is likely more sensitive to both mutational and environmental disruptions. Components of the cytosol may keep the highly mutated P7A7 somewhat functional, but extraction to an *in vitro* environment leads to a nonfunctional PTC. Of particular note are two chaperones, ObgE and EngA, that are proposed to interact with the PTC during assembly and may help direct PTC folding away from inactive intermediates, such as the non-native fold seen in P7A7 (**Figure 5**).^35,57,58^ Thus, due to an unstable PTC, P7A7 ribosomes are not suited for detailed *in vitro* study of improved β-amino acid polymerization reactions by the ribosome. Future efforts toward an understanding could include optimizing a cytosol-mimicking buffer for purification and analysis or mapping *in vivo* RNA structure of P7A7 ribosomes to assess whether it possesses a more stable PTC in cells.

The assembly issues with P7A7 suggest that future ribosome engineering efforts may need to consider three layers of optimization. First, the ribosome must be directed toward a novel function, be it antibiotic resistance, translation efficiency, or new polymerization chemistry. Secondly, the engineered ribosome must interact safely with the rest of the cell, avoiding dominant lethal phenotypes by inhibition of overall translation. Finally, ribosomes must productively assemble and remain stable when deployed for synthesis or study. These three goals may often be in conflict, with the necessary mutation of highly conserved rRNA bases towards new ribosome function threatening to derail cell survival or ribosome assembly. To solve part of this problem, much effort has gone into developing orthogonal ribosomes that do not interfere with normal translation. Mutations to anti-Shine-Dalgarno sequences^59–61^ and tethering of ribosomal subunits ^62,63^ have created a platform for powerful ribosome engineering, but there is as of yet no way to select for efficient ribosome assembly. Indeed, some of these engineered orthogonal ribosomes are similarly affected by assembly defects.^11^ Although P7A7 was the top hit in a screen for β-amino acid incorporation^9^ and there was little reason to believe it harbored significant assembly defects from initial *in vivo* assays, our results highlight that assembly problems as a result of aggressive mutations can go undetected in traditional cell-based experiments. Future efforts directed to incorporate assembly into selections for new ribosome function could open new opportunities for engineering robust translation systems for sequence-defined polymer synthesis.

## Methods

### Plasmids and Cloning

All PCR reactions were performed using a Q5 DNA polymerase kit (NEB) and manufacturer recommended concentrations of primers, dNTPs, and enzyme. Mach1 cells (ThermoFisher) were used for all rRNA plasmid cloning and expression. See Table S2 for primer sequences. Plasmids pKK3535-P7A7 (Tet^R^), encoding the P7A7 23S rRNA, and pLK35 ^64^ (Amp^R^) were used for cloning and rRNA expression. Plasmid pLK35-WT-MS2 was generated by whole-plasmid PCR amplification of pLK35 and blunt end ligation using primers h98_MS2_F/R (MS2 tag fragments in capitals). Plasmid pLK35-P7A7-MS2 was generated by Gibson assembly of a PCR fragment from pKK3535-P7A7 with primers P7A7_ex_F/R and a fragment from pLK35-WT-MS2 using the corresponding complementary Gibson primers. pKK3535-040329-MS2 was created via iterative whole-plasmid PCR and blunt end ligation using primers h98_MS2_F/R and then primers 040329_F/R.

### Polysome profiling and rRNA quantitation

Polysome profiling was performed as described^65^ using Mach1 cells expressing the desired 23S rRNA-expressing plasmid. Fractions were collected and extracted twice with 1 volume phenol-chloroform (pH 4.5), washed with 1 volume chloroform and precipitated with 1/10th vol. 3M sodium acetate (pH 5.2) and 2.5 vol. ethanol at −20°C overnight. Precipitated RNA was washed with 300 μL cold 70% ethanol and resuspended in water. Ribosomal RNA was reverse transcribed using Superscript III reverse transcriptase (ThermoFisher) with up to 2 μg of input rRNA and primer MS2_quant_R (Table S2). RNA was hydrolyzed with 10 M NaOH at 37 °C for 30 min and cDNA was precipitated as before with sodium acetate and ethanol. The cDNA pellet was washed with 70% ethanol and resuspended in water, and concentrations were normalized to 1 ng/μL. Quantification of the ratio of MS2-tagged rRNA to WT rRNA was done via semi-quantitative PCR with Q5 DNA polymerase (NEB). Primers MS2_quant_F/R bind outside the MS2 tagged region of 23S cDNA, generating a 146 base pair (bp) band for MS2-tagged molecules and a 114 bp band for WT. Each sample was amplified twice with varying input cDNA amounts or cycle numbers to verify linear PCR amplification. Most samples were optimized around 0.2 ng input cDNA and 8x-12x cycles for a 10 μL PCR reaction. PCR products were run on a 15% acrylamide Tris-Borate-EDTA minigel and stained with SybrSAFE DNA stain. Gel images were quantitated using ImageJ.^66^

### WT Ribosome Purification

WT 50S/30S ribosomal subunits were prepared as described previously^67^ except the 50S/30S peak was isolated alone and subjected to an additional spin over a 15-40% sucrose gradient in buffer C (20 mM Tris-HCl pH 7.5, 60 mM NH_4_CI, 6 mM MgCl_2_ 0.5 mM EDTA, 2 mM DTT). Fractions were concentrated and sucrose was removed using 100 kDa cutoff spin filters (millipore). The purified subunits were frozen in aliquots at −80 °C.

### P7A1 and 040329 Ribosome Crude Purification

*E. coli* Mach1 cells (ThermoFisher) were transformed with plasmid pLK35-P7A7-MS2 or pKK3535-040329-MS2. Overnight cultures of P7A7-MS2 expressing cells were diluted 1:100 into 3L LB + 100 μg/mL ampicillin (P7A7) or 50 μg/mL tetracycline (040329). P7A7 50S subunits were then prepared as described^67^ up through crude ribosome pelleting. The pellet was resuspended in ~2mL of buffer A (20 mM Tris-HCl pH 7.5, 100 mM NH_4_CI, 10 mM MgCl_2_ 0.5 mM EDTA, 2 mM DTT) and stored at 4 °C until MS2-tag affinity purification.

### MBP-MS2 purification

A pMAL-c2 plasmid (NEB) encoding N-terminally 6xHis tagged MBP-MS2 (Gift from Nadège Liaud, UC-Berkeley) was transformed into BL21 Codon+ RIL chemically competent cells (Agilent). An overnight culture was used to inoculate 3L of ZYM-5052 auto-inducing media (Studier) with 100 μg/mL ampicillin and grown for 16 h at 37 °C. Cells were cooled and all subsequent steps carried out at 4 °C. After pelleting at 4000 *g*, cells were washed and resuspended in 130 mL lysis buffer (20 mM HEPES pH 7.5, 250 mM KCl, 10 mM imidazole, 1 protease inhibitor tablet (Pierce) per 50 mL, 2 mM 2-mercaptoethanol) before lysis by sonication. Lysate was clarified by centrifugation (48,000 *g*, 30 min) and filtered through a 0.2 μM filter. MBP-MS2 was initially purified using an Akta fast protein liquid chromatography (FPLC) system (GE) with a 5 mL HisTrap column (GE). The column was washed with 10 column volumes (CV) of lysis buffer and eluted with a 10 CV linear gradient of lysis buffer with 10-500 mM imidazole. Protein containing fractions were combined and dialyzed against a low salt MS2 buffer (MS2-LS) (20 mM HEPES pH 7.5, 20 mM KCl, 1 mM EDTA, 2 mM 2-mercaptoethanol) in 10 kDa cutoff dialysis cassettes overnight. MBP-MS2 was then purified again on an FPLC using a 5 mL Heparin column (GE). Batches of 50-100 mg protein (estimated by A280, ε=83310 M^−1^cm^−1^) were bound to the column, washed with 5 CV of MS2-LS, and eluted with a linear gradient of MS2-LS with 20 mM-1 M KCl. MBP-MS2 fractions were combined, glycerol added to 10%, and stored in aliquots at −80 °C.

### MS2-tagged ribosome purification

All steps were performed at 4 °C. Crude ribosomes (>60 mg) were diluted to ~15 mg/mL in buffer A. MBP-MS2 (10 mg) was diluted to 0.5 mg/mL in MS2-150 buffer (20 mM HEPES pH 7.5, 150 mM KCl, 1 mM EDTA, 2 mM 2-mercaptoethanol) and loaded onto a 5 mL MBPtrap column (GE). The column was washed with 5 CV buffer A, loaded with crude ribosomes, and then attached to an FPLC. Ribosomes were purified with the following program: 5 CV wash with buffer A, 5 CV wash with buffer A with 250 mM NH_4_CI, 10 CV elution with a linear gradient of buffer A with 0-10 mM maltose. Ribosome containing fractions were concentrated and washed with buffer A in 100 kDa cutoff spin filters. Concentration was estimated with the conversion 1 nM 70S ribosomes = 24 A260. MS2-purified ribosomes were stored at −80 °C. For the short peptide *in vitro* translations, P7A7 ribosomes were additionally purified at small scale using the same method over ~100 μL (packed volume) amylose resin. Ribosomal purity was assayed using semi-quantitative RT-PCR as described in the polysome profiling assay.

### Nanoluciferase in vitro translation

*In vitro* translation (IVT) reactions used the PURExpress Δribosome kit (NEB) and a nanoluciferase reporter plasmid (Promega). IVTs used 0.5-1 μM purified ribosomal subunits, 10 ng/μL reporter plasmid, and all other components as per the manufacturer’s recommendation. Reactions (3 μL) were incubated for 60 min at 37 °C and quantified with the Nano-glo assay system (Promega).

### Short peptide translation

#### Formation of acyl-tRNAs used for protein synthesis

Synthesis, purification, and aminoacylation of tRNA^Val^ was carried out using the protocol described previously.^23^ Specifically, 10 μL of 250 μM Flexizyme (eFx)^68^ was added to 10 μL of either 200 mM HEPES with 200 mM KCl (pH 7.5, for α-Phenylalanine) or 500 mM HEPES (pH 7.5 for β-Phenylalanine) and 10 μL of 250 μM tRNA^Val^. The samples were incubated at 95 °C for 2 min and allowed to reach room temperature in 5 min. 60 μL of 1 M magnesium chloride was then added, followed by 10 μL of a DMSO solution of each cyanomethyl ester amino acid variant (50 mM). Reactions were incubated at 4 °C for 60 h. The reactions were quenched by addition of sodium acetate (pH 5.2) to a final concentration of 300 mM and ethanol was added to a final volume of 70% (*v/v*). The samples were then incubated at −80 °C for 1 h and the RNA was pelleted by centrifugation at 21,300 *g* for 30 min at 4 °C. The supernatant was removed and the pellet was washed with 500 μL of 70% (*v/v*) ethanol (stored at −20 °C). The sample was then centrifuged at 21,300 *g* for 7 min at 4 °C and the supernatant was removed. The pellet was air-dried for 2-5 min either at room temperature or on ice. When used immediately, the pellet was resuspended in 1 mM sodium acetate (pH 5.2). If used at a later date, the pellet was stored dry at −80 °C and resuspended in 1 mM sodium acetate (pH 5.2) before use.

#### In vitro translation reactions

Templates for *in vitro* translation of short peptides containing a FLAG tag fMet-Val-Phe-Asp-Tyr-Lys-Asp-Asp-Asp-Asp-Lys (MVFDYKDDDDK, fMVF-Flag) were generated as described previously.^23^ *In vitro* transcription/translation of (fMVF-Flag was carried out using the combination of PureExpress (ΔtRNA, Δaa (E6840S)) and PureExpress (Δribosome (E3313S)) kits (NEB) with the following modifications. To generate the fMVF-Flag WT peptide reactions contained (25 μL): Solution A (ΔtRNA, Δaa kit), 330 μM methionine, 330 μM valine, solution containing 330 μM tyrosine, 330 μM phenylalanine, and 330 μM lysine, 280 μM aspartic acid (pH 7), tRNA solution (ΔtRNA, Δaa kit), Factors Mix (Δribosome kit), 0.8 μM purified 30S subunits, 0.8 μM purified 50S subunits (either WT or P7A7), 500 ng dsDNA template, and water (to 25 μL). When using tRNA^Val^ (50 μM) charged using eFx, valine was omitted from the reaction mixture. The reactions were then incubated for 6 h at 37 °C. The reactions were quenched by placing them on ice and adding 25 μL of dilution buffer (10 mM magnesium acetate (Sigma-Aldrich) and 100 mM sodium chloride (Sigma Aldrich)). To remove the proteins and majority of nucleic acid macromolecules, 5 μL of Ni-NTA (Qiagen) slurry was added and the solution was incubated with light agitation at 4 °C for 50 min. The Ni-NTA resin was removed by centrifugation at 21,300 *g* for 10 min at 4 °C. The supernatant was then frozen at −80 °C for 5 min and centrifuged once more at 21,300 *g* for 10 min at 4 °C. The supernatant was analyzed on a Poroshell 120 EC-C18 column (2.7 μm, 3.0 × 50 mm, 45 °C, Agilent) using a linear gradient from 5 to 55% acetonitrile over 6.5 min with 0.1% formic acid as the aqueous mobile phase after an initial hold at 95% 0.1% formic acid for 0.5 min (0.6 mL/min) using a 1290 Infinity II UHPLC (G7120AR, Agilent). Peptides were identified using LC-HRMS as described previously.^23^

### Magnesium-dependent 70S complex formation

10 pmol WT or P7A7 50S subunits were mixed with 20 pmol WT 30S subunits in 20 μL of buffer C, described above, with 10 mM, 15 mM or 20 mM MgCl_2_ and incubated for 30 min at 37 °C. Reactions were spun over 15-30% sucrose gradients made in buffer C with appropriate MgCl_2_ concentrations at 178,000 *g* (SW-41 rotor, Beckman Coulter) for 3 hours. A254 traces were measured with an ISCO gradient fractionation system.

### EM sample preparation

Samples were deposited onto glow-discharged 300 mesh Quantifoil UltraAuFoil R1.2/1.3 grids with an additional top layer of continuous amorphous carbon floated on. The sample was initially incubated on the grid for approximately 1 minute, after which excess sample was washed off in a buffer containing 20 mM Tris-HCl, pH 7.5, 60 mM NH_4_Cl, 6 mM MgCl_2_ 0.5 mM EDTA, 2mM DTT. A Vitrobot Mark IV was used for plunge-freezing with the settings: 20 °C, 100% humidity, blot force 8, blot time 3 seconds. Inside the Vitrobot, the grid was first side-blotted with filter paper to remove the majority of the solvent, followed by depositing 1.2 μL of buffer prior to blotting and plunging into liquid ethane.

### EM Data Collection

Images were collected on an FEI Titan Krios electron microscope operated at 300 keV and with a GIF energy filter. The images were collected on a GATAN K2 Summit camera in super-resolution mode. Magnification was set to 215,000x for a pixel size 0.56 Å (0.28 Å super-resolution size). We collected images using a focal pair approach that we have since determined to have no advantage over a conventional defocus ramp. Briefly, two movies were recorded of each area, the first targeting a defocus level roughly around −0.3 μm (P7A7) or −0.5 μm (WT) with a total dose of 10 e-/ Å2, and the second with a target defocus around −3.5 μm (P7A7) or −3.6 μm (WT) and total dose 20 e-/ Å2. SerialEM^69^ was used for automated data collection and Focus^70^ was used for real-time monitoring of the data.

### Image processing

For all movies, motion correction was performed with MotionCor2.^71^ CTF estimation was performed with CTFFind4,^72^ and poorly fit micrographs were discarded based upon visual inspection. For ease of working with the focal pair data, far-focused movies were first processed to determine particle position and rough orientation before moving to the near-focused movies for high resolution structure determination.

For P7A7, templates for automatic picking were generated in RELION 2.0^33^ by first picking using a Gaussian blob, then classifying the results and choosing the best classes for templates. Three rounds of 2D classification were performed on 92,647 template-picked particles, reducing the number to 79,268 particles. Ab-initio reconstruction with 3 classes was performed in cryoSPARC v1,^34^ resulting in one class that resembled a normal 50S subunit and one class that appeared to be completely missing the central protuberance (CP). These two classes were separately submitted for 3D auto-refinement in RELION. The latter structure (with 21,319 particles from the far-focused data) did not refine to high enough resolution to pursue further. The class with an intact CP was subject to a round of 3D classification without alignment. Three out of four classes were pooled and refined to 4.17 Å from far-focused data. The resulting particle positions and orientations were then applied to near-focus movies with custom python scripts, and those particles were refined starting from local angular searches to 3.11 Å with 37,609 particles.

For WT 50S, all processing was done in RELION 2.0.^33^ 109,802 particles were auto-picked from far-focused micrographs with a Gaussian blob. Four rounds of 2D classification were performed to yield 99,356 particles for an initial 3D auto-refinement. Processing was shifted to near-focused micrographs as described for P7A7. 3D classification without alignment was performed after an initial near-focused refinement from local searches. The best class, containing 84,372 particles, was refined again for a final resolution of 3.20 Å. Coordinates for the WT 50S subunit based on PDB entry 4YBB^38^ were docked into the density as a rigid body. Molecular graphics were created using ChimeraX.^73^

### Cryo-EM map occupancy calculation

Map occupancy at each feature was calculated using a previously developed algorithm.^37^ A suitable contour level for each map was chosen after amplitude scaling and resampling by normalizing to the volume of uL4, a known early assembling ribosomal protein. Complete-linkage clustering was used to group structures by occupancy similarity. Results matched well with visual inspection of map features.

### LC-MS/MS and TMT-based quantification

#### Peptide preparation

Ribosomal proteins were precipitated in triplicate from WT-MS2 or P7A7-MS2 purified 50S subunits with 20% trichloroacetic acid at 4 °C for 1 hr. Protein pellets were washed 3x with 500 μL 0.01 M HCl in 90% acetone and dried. Protein was digested and TMT labeled using the TMTduplex isobaric mass tagging kit (ThermoFisher).

#### Mass spectrometry

Mass spectrometry was performed by the Vincent J. Coates Proteomics/Mass Spectrometry Laboratory at UC Berkeley. Peptides were analyzed on a ThermoFisher Orbitrap Fusion Lumos Tribrid mass spectrometry system equipped with an Easy nLC 1200 ultrahigh-pressure liquid chromatography system interfaced via a Nanospray Flex nanoelectrospray source. Samples were injected on a C18 reverse phase column (25 cm x 75 μm packed with ReprosilPur C18 AQ 1.9 μm particles). Peptides were separated by a gradient from 5 to 32% acetonitrile in 0.02% heptafluorobutyric acid over 120 min at a flow rate of 300 nL/min. Spectra were continuously acquired in a data-dependent manner throughout the gradient, acquiring a full scan in the Orbitrap (at 120,000 resolution with an AGC target of 400,000 and a maximum injection time of 50 ms) followed by 10 MS/MS scans on the most abundant ions in 3 s in the dual linear ion trap (turbo scan type with an intensity threshold of 5000, CID collision energy of 35%, AGC target of 10,000, maximum injection time of 30 ms, and isolation width of 0.7 m/z). Singly and unassigned charge states were rejected. Dynamic exclusion was enabled with a repeat count of 1, an exclusion duration of 20 s, and an exclusion mass width of ±10 ppm. Data was collected using the MS3 method^74^ for obtaining TMT tag ratios with MS3 scans collected in the orbitrap at a resolution of 60,000, HCD collision energy of 65% and a scan range of 100-500.

#### Data analysis

Protein identification and quantification were done with IntegratedProteomics Pipeline (IP2, Integrated Proteomics Applications, Inc. San Diego, CA) using ProLuCID/Sequest, DTASelect2 and Census [3,4,5,6],^75–77^ Tandem mass spectra were extracted into ms1, ms2 and ms3 files from raw files using RawExtractor^78^ and were searched against the *E. coli* protein database plus sequences of common contaminants, concatenated to a decoy database in which the sequence for each entry in the original database was reversed.^79^ All searches were parallelized and searched on the VJC proteomics cluster. Search space included all fully tryptic peptide candidates with no missed cleavage restrictions. Carbamidomethylation (+57.02146) of cysteine was considered a static modification; TMT tag masses, as given in the TMT kit product sheet, were also considered static modifications. We required 1 peptide per protein and both tryptic termini for each peptide identification. The ProLuCID search results were assembled and filtered using the DTASelect^76^ program with a peptide false discovery rate (FDR) of 0.001 for single peptides and a peptide FDR of 0.005 for additional peptide s for the same protein. Under such filtering conditions, the estimated false discovery rate was zero for the dataset used. Quantitative analysis on MS3-based MultiNotch TMT data was analyzed with Census 2 in IP2 platform.^80^ As TMT reagents are not 100% pure, we referred to the ThermoFisher Scientific TMT product data sheet to obtain purity values for each tag and normalized reporter ion intensities. While identification reports best hit for each peptide, Census extracted all PSMs that can be harnessed to increase accuracy from reporter ion intensity variance. Extracted reporter ions were further normalized by using total intensity in each channel to correct sample amount error.

To account for purity differences between ribosomal protein samples, two additional normalization methods were attempted. First, intensities were normalized to the total 50S ribosomal protein signal for each respective mass tag. Second, intensities were normalized to the averaged signals of early assembling proteins uL4, uL13, bL20, uL22, and uL24. These proteins are expected to be be stoichiometrically present in both mature WT 50S and assembly-stalled P7A7 subunits. These two methods produced similar results and the latter was chosen for further analysis. Reported proteins were identified in all three relative quantitation experiments with the exception of L35, which was seen in two.

## Supplementary Figures

**Figure S1.**
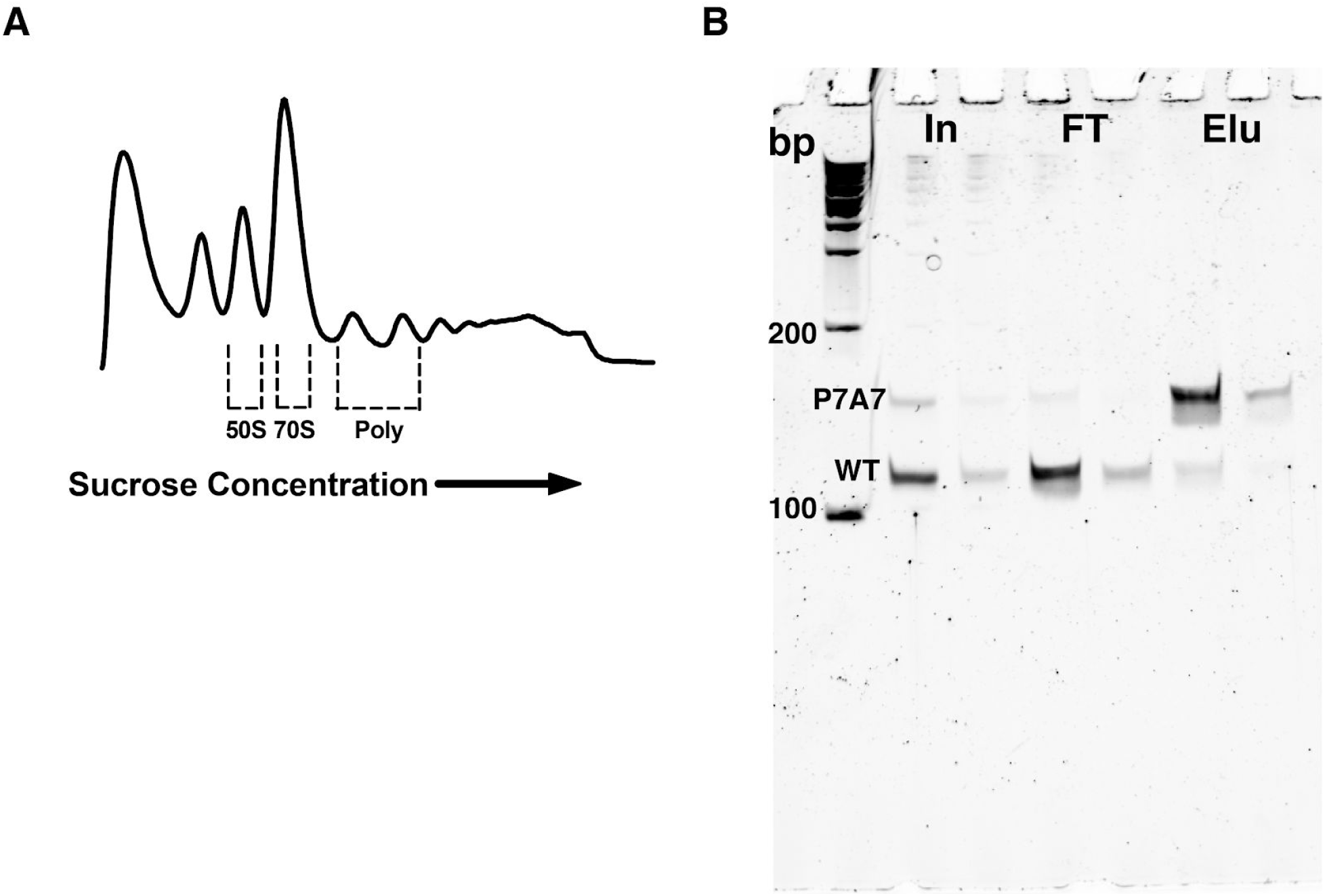
Distribution of P7A7 50S subunits in polysomes. A) Representative sucrose gradient fractionation of polysomes showing fractions collected for analysis in **Figure 1A.** The 254 nm absorbance trace is shown. B) P7A7 purification assayed by semi-quantitative RT-PCR over the MS2-tag of isolated rRNA. Input (In), flow through (FT), and elution (Elu) fractions analyzed via two PCRs for each sample, with 0.2 ng and 0.04 ng of input cDNA for the stronger and weaker lanes, respectively.

**Figure S2.**
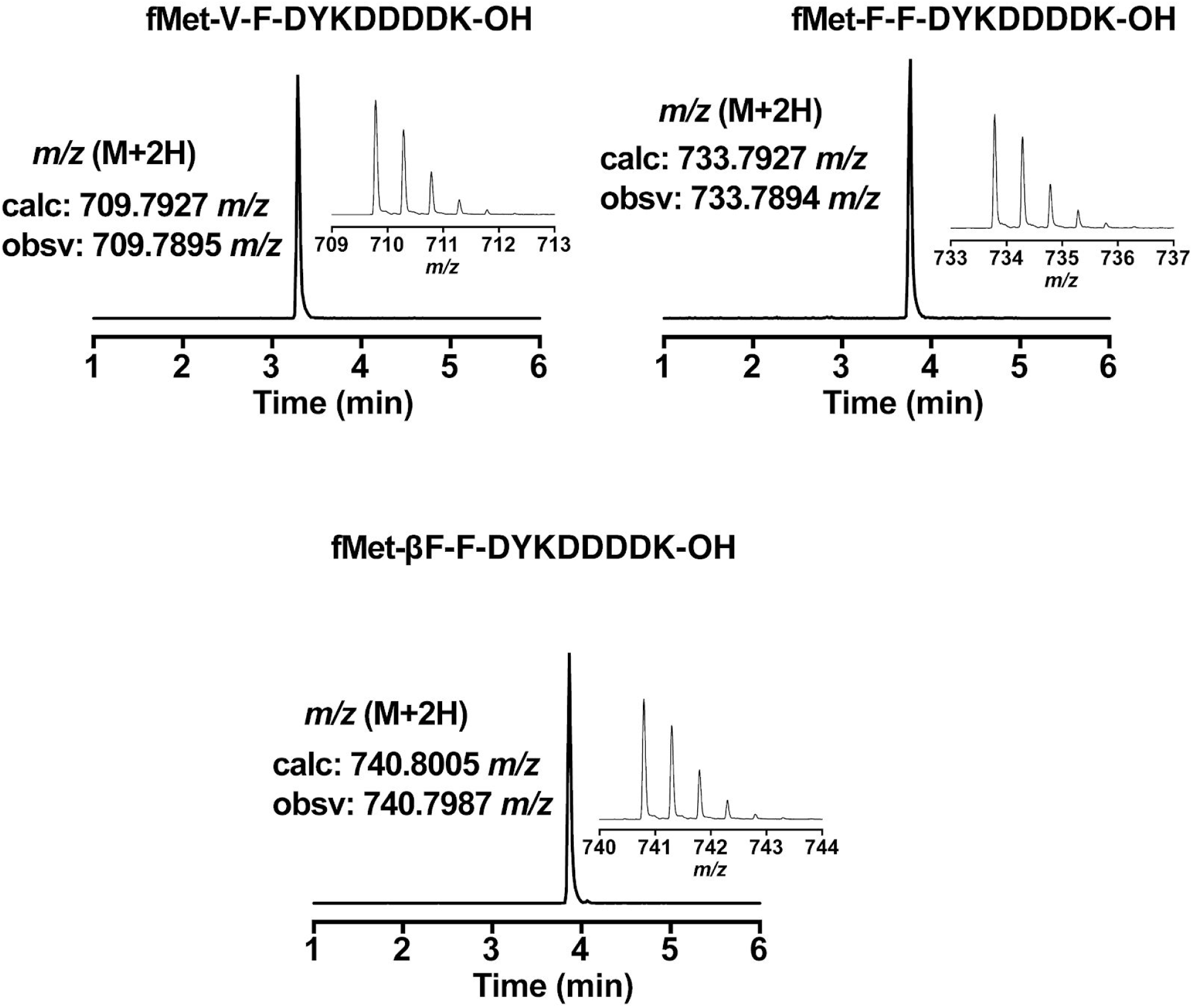
Extracted ion chromatograms and mass spectra of FLAG-derived peptides. Liquid chromatography and mass spectrometry results for the *in vitro* translation experiments in **Figure 1C**.

**Figure S3.**
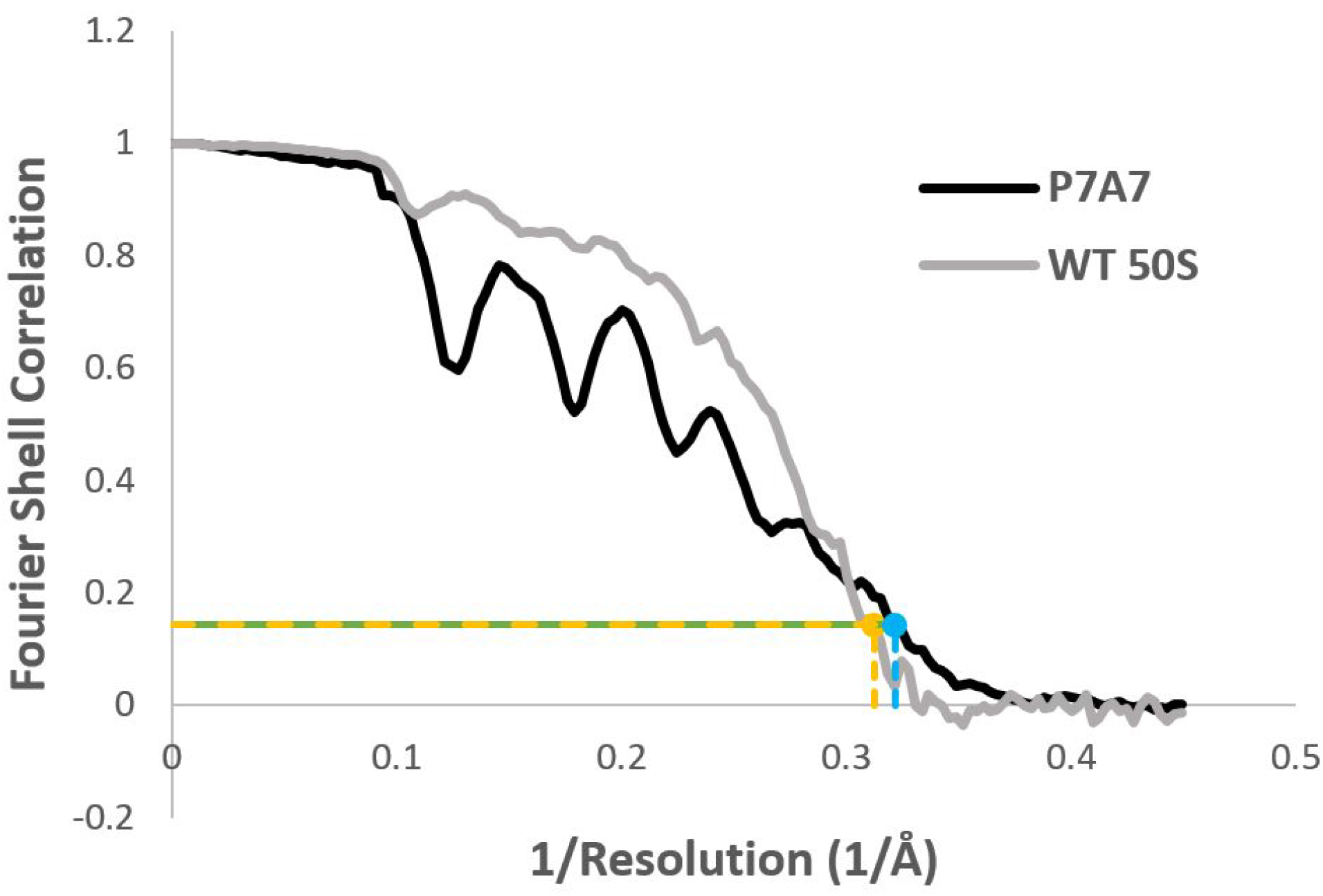
Fourier shell correlation (FSC) curves for P7A7 and WT 50S cryo-EM maps. Oscillations in the FSC for P7A7 are an artifact of the focal pair method of data collection, caused by zeroes in the CTF at the predominant defocus value. The same effect is not observed for the WT 50S subunit map due to different targeting of the near-focus value during the data collection session. The gold-standard FSC cutoff value for global resolution (0.143) is marked in gold for WT and blue for P7A7.

**Figure S4.**
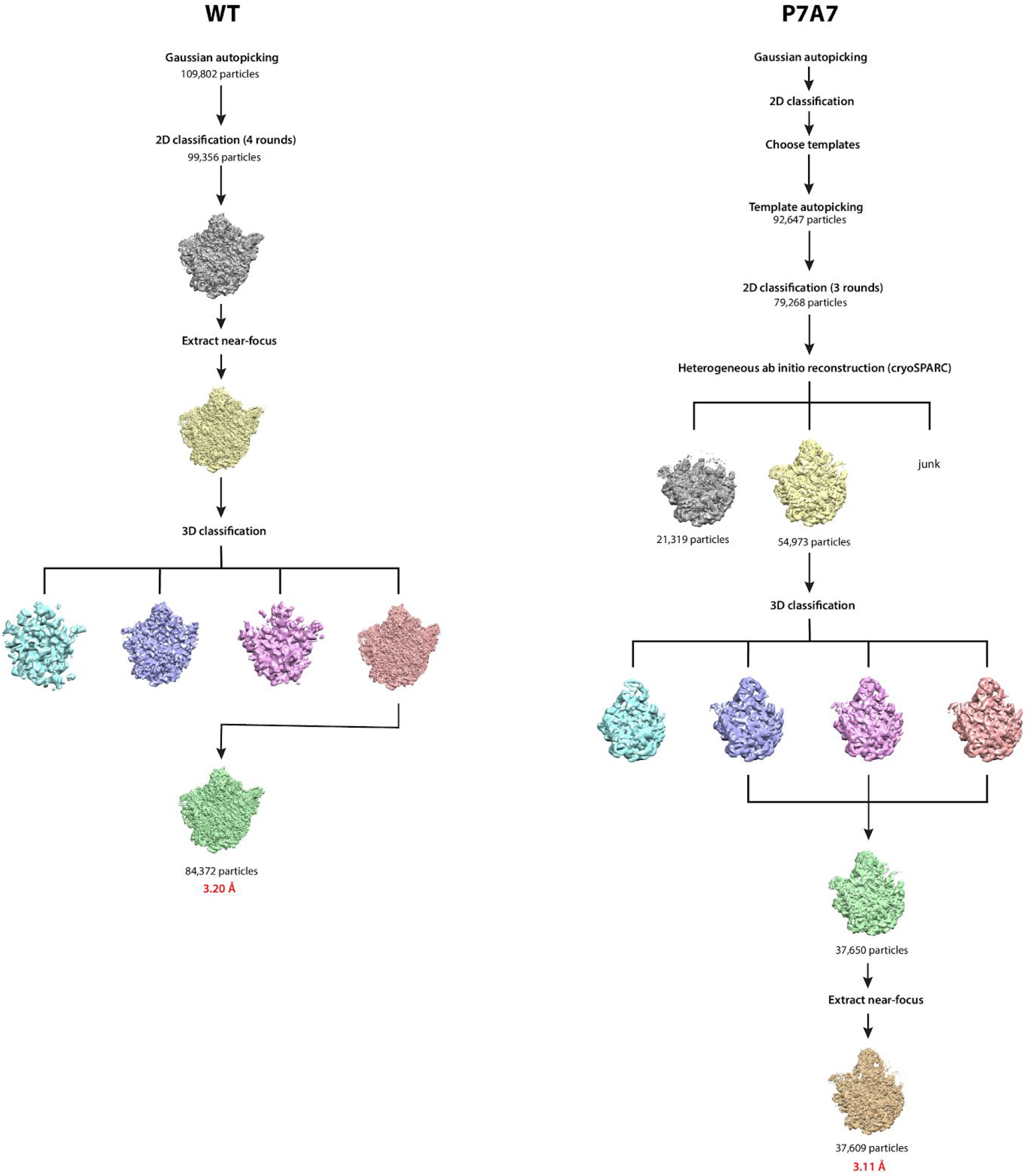
Cryo-EM data processing workflow for P7A7 and WT 50S subunits. Most steps were completed in RELION, otherwise in cryoSPARC where noted.The smaller class of P7A7 subunits lacking the CP is shown in grey.

**Table S1.** Cryo-EM data collection and refinement information.

|  | P7A7 large subunit | WT 50S |
| --- | --- | --- |
| Microscope | FEI Titan Krios |  |
| Accelerating Voltage (kV) | 300 |  |
| Detector | GATAN K2 Summit |  |
| Spherical Aberration (mm) | 2.7 |  |
| Magnification | 215,000x |  |
| Defocus targets ($\mu\text{m}$) | -0.3 and -3.5 | -0.5 and -3.6 |
| Micrographs (defocus pairs) | 1402 | 1442 |
| Particles picked | 92,647 | 109,802 |
| Particles refined | 37,609 | 84,372 |
| Resolution achieved ($\text{\AA}$) | 3.11 | 3.20 |

**Table S2:**
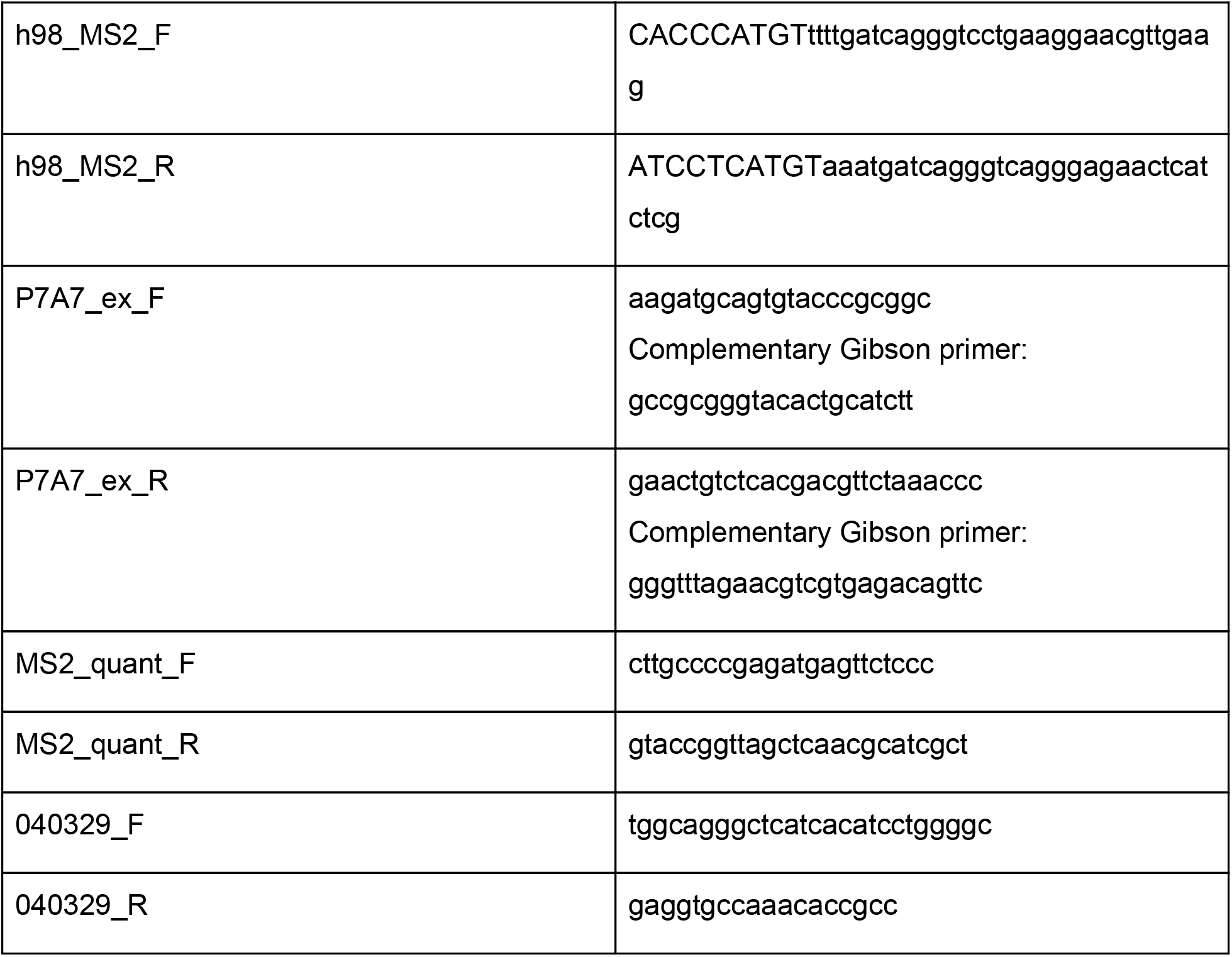
Primers used in this study.

## Acknowledgements

We thank Bong-Gyoon Han, Dan Toso, and Paul Tobias for assistance with cryo-EM sample preparation and data collection, as well as Lori Kohlstaedt and Carl Ward for mass spectrometry advice. Mass spectrometry was performed by the Vincent J. Coates Proteomics/Mass Spectrometry Laboratory, QB3 Institute, UC Berkeley. The cryo-EM work reported herein was supported by the Center for Genetically Encoded Materials, an NSF Center for Chemical Innovation (NSF CHE-1740549) and by an NSF predoctoral fellowship to Z.W. (1106400). The biochemical studies were supported by the NIH (GM R01-114454). O.A. was supported in part by Agilent Technologies as an Agilent Fellow.

